# Extracellular electron transfer drives efficient H_2_-independent methylotrophic methanogenesis by *Methanomassiliicoccus,* a seventh order methanogen

**DOI:** 10.1101/2024.03.29.587405

**Authors:** Lingyan Li, Xuping Tian, Xuemeng Wang, Chuan Chen, Qi Zhou, Lei Qi, Jie Li, Kai Xue, Fangjie Zhao, Yanfen Wang, Xiuzhu Dong

## Abstract

Methylotrophic methanogenesis is achieved via methyl group dismutation or H_2_ reduction. This study reports extracellular electron droving efficient methylotrophic methanogenesis. The 7^th^ order methanogen *Methanomassiliicoccus luminyensis* exclusively implements H_2_-dependent methylotrophic methanogenesis, but strain CZDD1 isolated from paddy soil possessed a higher methane-producing rate in coculture with *Clostridium malenominatum* CZB5 or the electrogenic *Geobacter metallireducens.* Chronoamperometry detected current production from CZB5, and current consumption accompanied CH_4_ production in a methanol-containing electrochemical culture of CZDD1. This demonstrated that *M. luminyensis* was capable of both direct species electron transfer (DIET) and extracellular electron transfer (EET) in methylotrophic methanogenesis. EET and DIET also enabled CZDD1 to produce methane from dimethyl arsenate. Differential transcriptomic analysis on H_2_-versus EET- and DIET-cocultures suggested that a membrane-bound Fpo-like complex and archaella of *M. luminyensis* CZDD1 could accept extracellular electrons. Given the ubiquitous environmental distribution of *Methanomassiliicoccus* strains, EET driven methylotrophic methanogenesis may contribute significantly to methane emission.

## Introduction

Being the second most significant greenhouse gas after carbon dioxide (CO_2_), methane (CH_4_) contributes to approximately 30% of global warming. Biogenic methane formed through methanogenesis accounts for approximately 70% of atmospheric CH_4_ emissions, and methanogenic archaea are the only known organism producing ample methane. Methanogenic archaea produce CH_4_ through three main pathways: CO_2_ reductive or hydrogenotrophic, aceticlastic, and methylotrophic methanogenesis, and the latter pathway includes two subtypes, H_2_-dependent-methyl group reduction, and methyl group dismutation or disproportionantion (*1*). Recently, electromethanogenesis based on direct interspecies electron transfer (DIET) has been reported, such as DIET between *Geobacter metallireducens* and *Methanosarcina barkeri*, and *Methanosarcina acetivorans*, in which outer-surface multiheme c-type cytochrome takes up the extracellular electrons (*2, 3*). While DIET facilitated methanogenesis was also reported in cytochrome C lacking Methanobacterium (*4*). DIET even activated the CO_2_ reductive methanogenic pathway encoded by *Methanotrix* (*Methanosaeta*) *harundinacea*, an obligately aceticlastic methanogen (*5*). *Geobacter* species are the pioneering bacteria found to donate electrons to many partners (*6*). Later, electrodes were found to be capable of extracellular electron transfer (EET) to numerous bacteria, such as *Clostridium* spp. and *Listeria monocytogenes* (*6–8*). Extracellular electron transfer (EET) from cathode was reported to support methane production by *Methanococcus maripaludis* and *Methanosarcina* spp. (*9*). This suggests that DIET and EET maybe commonly used by methanogens.

Recent metagenomic investigations have indicated that the H_2_-dependent methylotrophic methanogenic pathway is present in multi-archaeal clades including five orders within the Euryarchaeota phylum, and the phyla of Ca. Verstraetearchaeota and Korarchaeota (*10, 11*). Moreover, this pathway is widely distributed in diverse environments (*12–17*), suggesting that its ecological significance has been underestimated. Methanomassiliicoccales methanogens, the so-called 7^th^ order methanogens, belong to the Thermoplasmata phylum and grow exclusively on H_2_-dependent methylotrophic methanogenesis. They are widely distributed in various environments, including ocean (*18*), rice paddy soil (*19*), mammalian gut (*20–22*), mangrove sediments (*23*), and freshwater wetlands (*24*). However, *Methanomassiliicoccus luminyensis* B10 is one of the few cultured Methanomassiliicoccales strains and firstly isolated from the human intestine (*25*). It slowly grows on H_2_-dependent methanol or trimethylamine derived methanogenesis (doubling times of 1.8 d and 2.1-2.3 d)(*26*). The genome of strain B10 does not encode cytochromes, and the encoded energy-converting hydrogenase is not inactive, therefore suggesting that it may generate proton motive force (PMF) for ATP formation by a membrane-bound Fpo-like complex coupled with redox of ferredoxin (*27*). Biochemical assays demonstrated ferredoxin oxidation coupled to heterodisulfide reduction by the cell membrane preparation of *M. luminyensis* B10 (*28*), providing further evidence for a mode of energy metabolism distinct from that of the classical methanogens.

Recently, we isolated another *M. luminyensis* strain CZDD1 from dimethylarsenate (DMAs)-contaminated paddy soil. Strain CZDD1 produced methane from methanol and trimethylamine using H_2_ as the reductant, and also produced methane from DMAs only in coculture with a co-isolated bacterium *Clostridium malenominatum* CZB5 (*29*). Remarkably, the coculture possessed a higher methane producing rate from methanol than the H_2_-dependent monoculture, implying that *C. malenominatum* CZB5 might provide other reductants, like extracellular electrons, than H_2_ to *M. luminyensis* CZDD1. Although DIET supported CO_2_ reductive methane production has been reported (*5, 9*), this activity remained unknown for methylotrophic methanogenesis. This work explored the potential of DIET and EET in methylotrophic methanogenesis by *M. luminyensis* CZDD1 based on physiological and electrochemical experiments. These demonstrated that DIET with *C. malenominatum* CZB5 and EET supported higher rates of methanogenesis by *M. luminyensis* CZDD1 than H_2_. Importantly, both DIET and EET but not H_2_ enabled methanogenesis from DMAs, a toxic chemical that causes straight-head rice disease (*30*). Based on differential transcriptomic analysis, the EET-based methanogenic pathway and energy metabolism mode was explored. This work extends knowledge of DIET and EET-based methanogenic pathways, which could contribute significantly to atmospheric methane evolution.

## Results

### *C. malenominatum* CZB5 enables *M. luminyensis* CZDD1 to produce methane in the absence of H_2_

Like *M. luminyensis* B10, pure culture of *M. luminyensis* CZDD1 grows and obtains energy exclusively via H_2_-dependent methylotrophic methanogenesis. Upon inoculating 10% of the *M. luminyensis* CZDD1 culture (about 0.1 mg total cell protein) into a fresh medium containing methanol and H_2_, methane production started after 7-day with a production rate of 23.5 μmol per day (Fig. 1A). However, when inoculating the same amount of *M. luminyensis* CZDD1 cells with a culture of *C. malenominatum* CZB5, methane production started after one day and obtained 1.9-fold higher rate of methanogenesis (43.75 μmol per day) in the absence of added H_2_ (Fig. 1A). H_2_ accumulated in both the *C. malenominatum* CZB5 monoculture and its co-culture with *M. luminyensis* CZDD1, but H_2_ disappeared as CH_4_ was produced in the coculture (Fig. 1B). In total, the coculture produced more than 300 μmol CH_4_ but only consumed about 18 μmol H_2._ This ratio was far lower than the 1:1 stoichiometry of H_2_ to CH_4_ expected in H_2_-dependent methylotrophic methanogenesis, suggesting that other reductants from CZB5 provided to *M. luminyensis* CZDD1. To further preclude the involvement of H_2_ in methanogenesis, a gas mixture of 10% carbon monoxide (CO) and 90% N_2_ at 0.2 MPa was added to both monoculture and coculture to inhibit NiFe hydrogenases. As *M. luminyensis* CZDD1 encodes both membrane-bound (RS01470-RS01475, RS15345-RS15350, RS12290-RS122305) and cytoplasmic Ni-Fe hydrogenases (RS07015 and RS07025). *C. malenominatum* CZB5 also encodes a number of Ni-Fe hydrogenases (GM000120, GM000623, GM000648, GM000996, GM001621, GM001728, GM001731, GM001840, GM002037, GM002260, GM002441, GM002630, GM002892, GM002962, GM003036 and GM003561). Amendment of CO abolished methane production in *M. luminyensis* CZDD1 monoculture, but only reduced methanogenic rate of the coculture to 30.05 μmol per day (Fig. 1A), which was still higher than the rate of methane production by the CZDD1 monoculture. CO significantly reduced H_2_ production by *C. malenominatum* CZB5 monoculture (Fig. 1B), but not the growth (Fig. S1).

**Figure 1.**
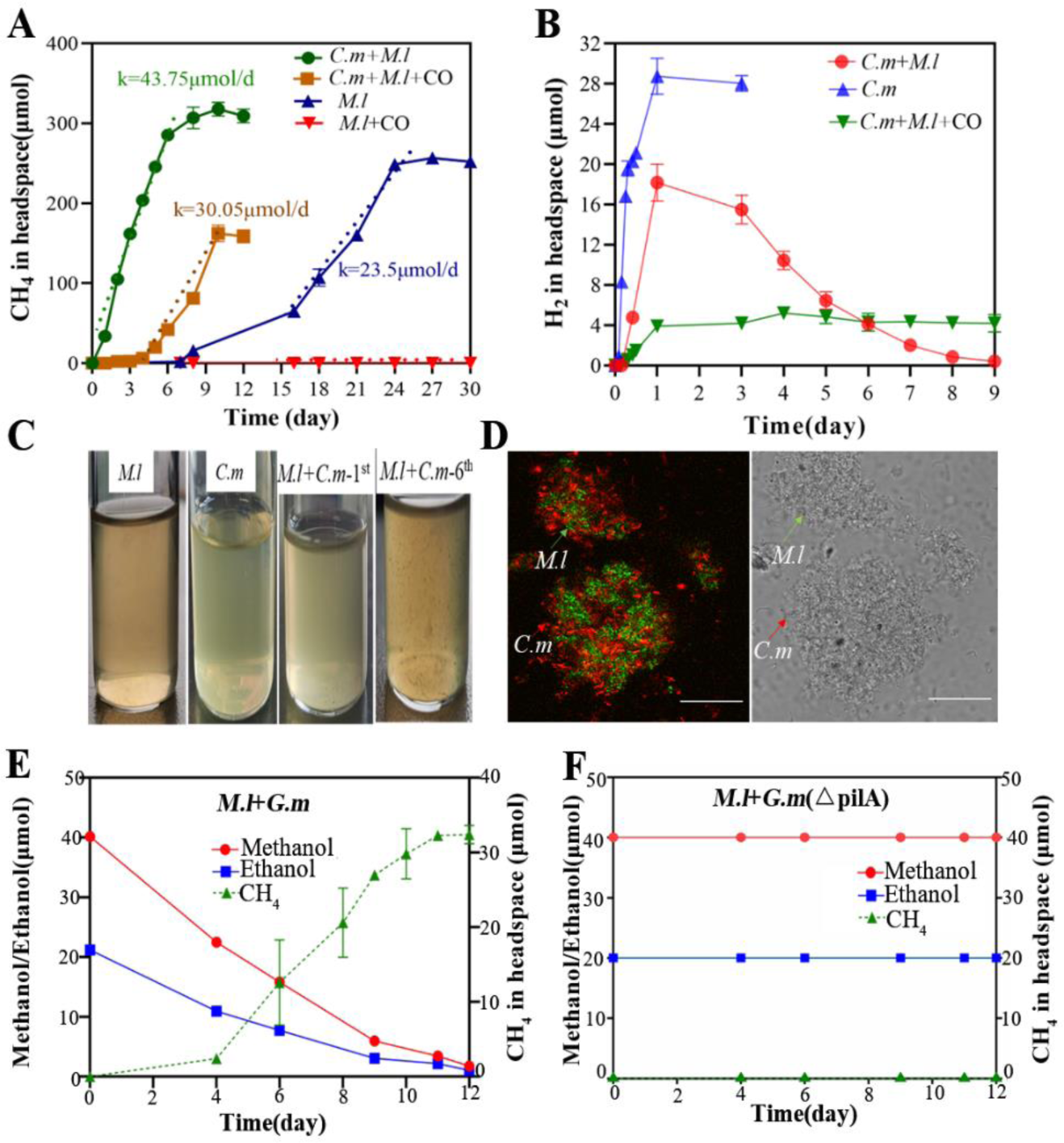
DIET facilitates efficient H_2_-independent methylotrophic methanogenesis by *M. luminyensis*. Pre-reduced modified MM medium was used in all experiments, and all the cultures were initiated with the same inoculation sizes and incubated at 37 °C. (**A** and **B**) *M. luminyensis* CZDD1 (*M.l*), *C. malenominatum* CZB5 (*C.m*) monocultures, and coculture of the two were all cultured in 375 μmol methanol. The *M. luminyensis* monoculture was purged with 0.1 MPa H_2_, and the *C. malenominatum* monoculture and the co-culture were purged with 0.1 MPa N_2_. 0.2 MPa of 10% CO and 90% N_2_ was added to cultures to inhibit [NiFe] hydrogenase activity. Methane (**A**) and H_2_ accumulations (**B**) were monitored. (**C**) Aggregations in the 6^th^ subcultures of the coculture. (**D**) Using Cy3 (red)- and Cy5 (green)-labeled 16S rRNAs to specifically probe *M. luminyensis* and *C. malenominatum*, respectively, FISH was performed on the coculture aggregates (left). Bright field micrograph of the same aggregates (right). Bars indicated 25 μm. (**E** and **F**) Cocultures were initiated with 10% each of *M. luminyensis* (*M.l*) and *G. metallireducens* (*G.m*, **E**), or its *pilA* deletion mutant (*ΔpilA*, **F**) into 200 μmol methanol and 100 μmol ethanol. Methane in headspace, and methanol and ethanol in liquid culture were monitored. Data are averages and standard deviations of three replicates.

To identify the possible reductants other than H_2_ transferred from *C. malenominatum* CZB5, its spent culture and the cell lysate were added to the *M. luminyensis* CZDD1 monoculture. However, none of the amendments enhanced the H_2_-dependent methylotrophic methanogenesis (Fig. S2). Thus, it seemed possible that the enhanced methanogenesis resulted from cell-to-cell contact. Indeed, although cell aggregates were not apparent in young cocultures, visible aggregates with diameters ranging 100 – 200 µm were observed after 6 generations of the coculture, but not in the monoculture (Fig. 1C). Using Cy3- and Cy5-labeled *M. luminyensis-* and *C. malenominatum-* specific 16S rRNA fragments as the respective probes, fluorescence in situ hybridization (FISH) determined roughly similar cell numbers of the two types of cells in the aggregates (Fig. 1D). These experimental results suggested that DIET was involved in methanogenesis in the coculture of *C. malenominatum* and *M. luminyensis*.

### DIET from *Geobacter metallireducens* enables H_2_-independent methylotrophic methanogenesis by *M. luminyensis* CZDD1

To test DIET involvement in H_2_-independent methylotrophic methanogenesis by *M. luminyensis* CZDD1, *Geobacter metallireducens*, a bacterium with a defined DIET activity, was cocultured with *M. luminyensis* CZDD1. *G. metallireducens* could not oxidize ethanol to acetate, protons, and electron by itself (formula 1), and *M. luminyensis* could not metabolize ethanol and methanol to methane without a reductant. If *M. luminyensis* could accept extracellular electrons from *G. metallireducens*, the coculture could reduce methanol to methane with ethanol as electron donor (formula 2). A coculture was constructed by 10% inoculation of each strain, and upon a 12-day incubation it consumed 100 μmol ethanol and 200 μmol methanol, and produced 160 μmol methane but no hydrogen (Fig. 1E), thus, about 80% of electrons from ethanol oxidation were channeled to methane production. However, no methane was detected in the co-culture of *M. luminyensis* CZDD1 with the deletion mutant of the *G. metallireducens pilA* (Fig. 1F), the gene that encodes e-pili for transferring extracellular electrons. This demonstrates that *M. luminyensis* CZDD1 has capacity of using DIET for methylotrophic methanogenesis. Like *M. luminyensis* CZDD1 in co-culture with *C. malenominatum* CZB5, its coculture with *G. metallireducens* also initiated methane production one day after inoculation (Fig. 1E), and suggesting that DIET might be preferred to H_2_ as an electron donor.

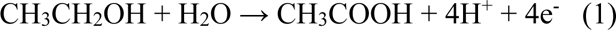

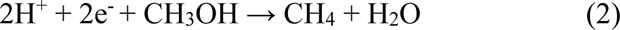

### Extracellular electron transfer by *C. malenominatum* CZB5

To determine the EET activity of *C. malenominatum* CZB5, its cultures were first amended with ferric citrate and ferrous iron production was measured. With 7.5 mM ferric citrate, Fe (III) reduction rate was 13.3 µM/min (Fig. 2A), approximately half of reduction rate (30 µM/min) of *Listeria monocytogenes* (*8*). Furthermore, during growth in an electrochemical reactor, *C. malenominatum* CZB5 yielded current with a density of approximately 20 µA/cm^2^ (Fig. 2B), similarly to that produced by *Listeria monocytogenes.* But no current was measured in the reactor without inoculation of the bacterium. These experiments confirm the extracellular electron transfer activity of *C. malenominatum* CZB5.

**Figure 2.**
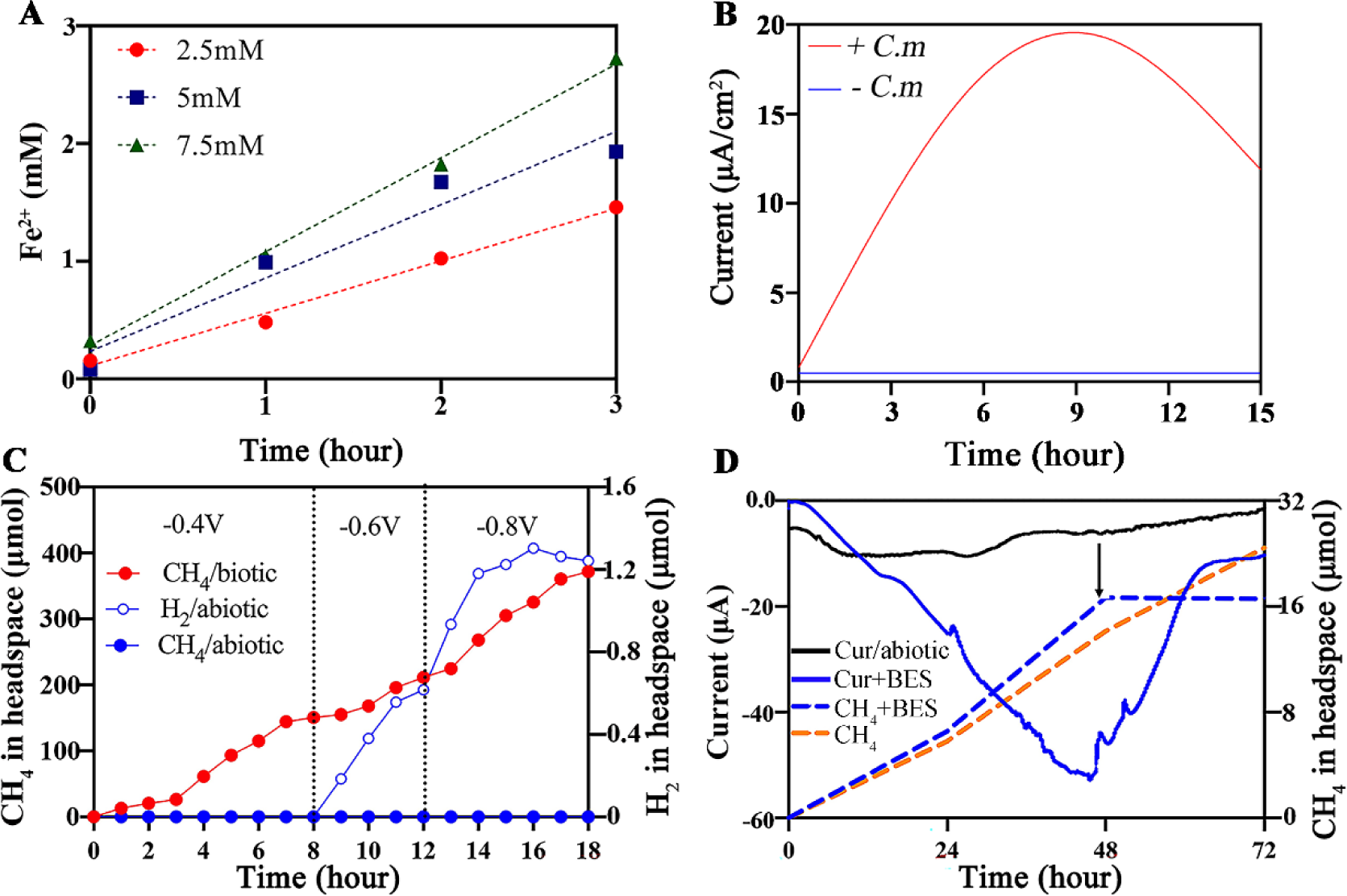
Extracellular electron transfers of *C. malenominatum*, and *M. luminyensis* in methylotrophic methanogenesis. (**A**) Reduction of ferric citrate by *C. malenominatum*. MM media containing the indicated concentrations of ferric citrate were given a 10% inoculation and incubated at 37°C. (**B**) MM medium without sulfide and resazurin in a cathode H-cell reactor with a potential of +0.4 V versus Ag/AgCl was given a 10% inoculation with *C. malenominatum* (*C.m*). An electrochemical culture without inoculation (-) served as abiotic control. The electrochemical cultures were mixed by a magnetic stir bar, and current production was monitored during incubation at 37°C. Experiments were performed on three batches of culture, and one representative dataset is shown. (**C**) A H-cell containing pre-reduced MM medium with 400 μmol methanol was given 10% inoculation with *M. luminyensis* (0.1 mg total cell protein), and purged with 100% N_2_ gas. An uninoculated cell served as an abiotic control. The cathode potential was set at −0.4 V in the first 8-day incubation at 37°C and then reduced to −0.6 V and −0.8 V as indicated. Productions of CH_4_ and H_2_ were monitored. (**D**) Current consumption and methane production were measured in the same electrochemical reactor as in **C** at −0.4 V potential of the cathode. A final concentration of 10 mM 2-Bromoethanesulfonate (BES) was added after 48 h as indicated by the arrow. Two independent experiments were performed, and one representative is shown here.

To determine the potential pathways of *C. malenominatum* CZB5 in extracellular electron transfer, transcriptomes were compared between its mono- and co-culture with *M. luminyensis* CZDD1 (Supplementary dataset S1). *C. malenominatum* CZB5 contains two suites of genes (GM002851-GM002852; GM002969-GM002970) encoding the electron transfer flavoprotein. Expression of GM002851-GM002852 increased 32-fold in the coculture compared with the monoculture, suggesting that FAD could act as an extracellular electron vector. In addition, expression of the genes encoding flagella and the chemotaxis system were also significantly upregulated (*P* < 0.05) in the coculture, suggesting that *C. malenominatum* CZB5 could move actively toward *M. luminyensis* CZDD1, and facilitate cell aggregation and direct interspecies electron transfer.

### EET enables methylotrophic methanogenesis by *M. luminyensis* CZDD1

During incubation of an early stationary phase culture of *M. luminyensis* CZDD1 in an electrochemical cell set at −0.4 V of a cathode versus the standard hydrogen electrode, a condition in which abiotic H_2_ would not be generated, methane production was detected at day one and achieved a similar production rate (29.7 µmol·d ^-1^) as the CO-amended coculture. Further lowering the cathode potential to −0.8 V increased the methanogenic rate to 40.2 μmol·d^-1^ (Fig. 2C). Abiotic H_2_ was detected at the cathode potential of −0.8 V but not at −0.4 V (Fig. 2D). This result suggested that EET directly enable methylotrophic methanogenesis by *M. luminyensis* CZDD1 without a H_2_ intermediate.

To verify the capability of *M. luminyensis* CZDD1 using cathodic electrons, methane production coupled to current consumption was assayed. At a cathode potential of −0.4 V, *M. luminyensis* CZDD1 consumed a current of about 40 μA and produced 14.2 μmol CH_4_ after 48 h. The addition of bromoethane sulfonate (BES), a specific inhibitor of the key methanogenic enzyme methyl-CoM reductase, simultaneously inhibited methane production and current consumption (Fig. 3D). This experiment demonstrated that *M. luminyensis* CZDD1 can use cathodic electrons to reduce methanol for methane production, i.e. performing an electromethanogenesis. Similar but lower activity was detected with *M. luminyensis* B10, a strain isolated from human intestine (Fig. S3), indicating that electromethanogenesis is a shared characteristic of the other strains of *M. luminyensis*.

**Figure 3.**
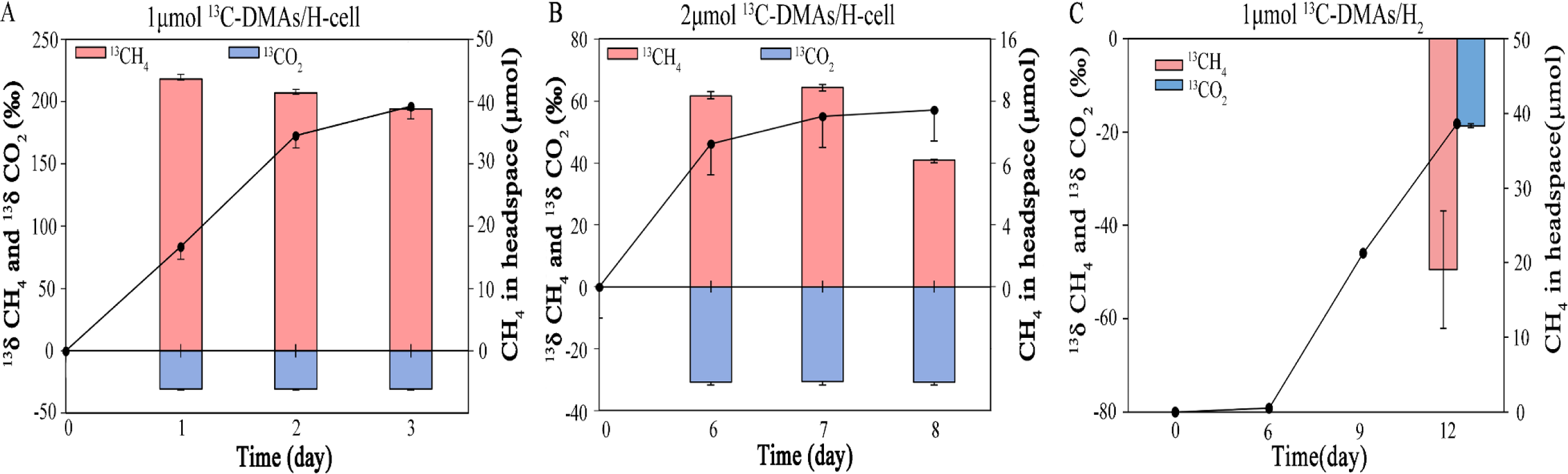
EET facilitates methane production from DMAs by *M. luminyensis*. (**A** and **B**) *M. luminyensis* CZDD1 was respectively cultured 400 μmol methanol amended with 1 μM (**A**) and 2 μM (**B**) of ^13^C-DMAs (V) and cultured in a H-cell reactor at −0.4 V of the cathode potential. he headspace contained 0.1 MPa N_2_. (**C**) A H_2_-culture containing 1 μM ^13^C-DMAs (V) and 400 μmol methanol was purged with 0.1 MPa H_2_. ll cultures were incubation at 37°C. Methane (black dot and line) was determined by gas chromatography, and ^13^CH_4_ (red bar) and ^13^CO_2_ (blue r) in the headspace were determined using GC - IRMS. Triplicate experiments were performed, and the means and standard deviations are shown.

### EET enables methanogenesis from DMAs(V) by *M. luminyensis* CZDD1

Recently, we reported that a *M. luminyensis* CZDD1 coculture with C. *malenominatum* CZB5 produced CH_4_ from DMAs (V) but the monoculture did not (*29*). It was also found that *C. malenominatum* CZB5 reduced DMAs (V) to DMAs (III), which is a highly toxic (*30*). Given that EET promoted a high rate of methanogenesis by *M. luminyensis* CZDD1 (Fig. 1E and 2C), we then explored whether EET could facilitate CH_4_ production from DMAs (V). Because of the toxicity of DMAs (III), low concentrations of DMAs(V) were used. *M. luminyensis* CZDD1 was incubated with 1 μM or 2 μM ^13^C labeled DMAs(V) mixed with 15 mM methanol in electrochemical chambers at −0.4 V of the cathode potential as above. After a few days of incubation, produced methane was ^13^C-enriched, in contrast, CO_2_ was not ^13^C-enriched and contained only the natural abundance (Fig. 3). For comparison, ^13^CH_4_ was not detected in the H_2_ culture of *M. luminyensis* CZDD1 containing 1 μM DMAs(V) and 15 mM methanol. Therefore *M. luminyensis* CZDD1 utilized cathodic electrons instead of H_2_ to produce CH_4_ from DMAs (V). Notably, a higher proportion of ^13^CH_4_ (average 200 ‰ vs. 60 ‰) was detected from the culture containing 1 μM ^13^C-DMAs (V) than that of 2 μM ^13^C-DMAs(V). This difference could be attributed to inhibition of *M. luminyensis* by DMAs (III) from DMAs (V) reduction. Whereas the CO_2_ stable carbon isotope signature was maintained at −30 ‰ in ^13^C-DMAs(V) amended culture (Fig. 3), indicating that the methyl group from DMAs was completely channeled to CH_4_, consistent with the absence of the genes encoding the methyl-group oxidization pathway of *M. luminyensis* CZDD1.

### Differential expressed *M. luminyensis* CZDD1 genes during DIET and EET cultures

To obtain the gene expression profiles of *M. luminyensis* CZDD1, transcriptomes were compared between H_2_-dependent and EET-dependent monocultures and the DIET coculture. Comparing with the H_2_-culture, expression of genes encoding the protein complexes potentially involved in the electron flow inside *M. luminyensis* CZDD1 were elevated in the coculture (Fig. 4A), that include *fpoABCDHIJKLMN* encoding a Fpo-like complex, and *hdrABC*/*mvhAGD* encoding cytoplasmic heterodisulfide reductase-hydrogenase complex, and *hdrD* encoding a subunit of membrane-bound heterodisulfide reductase HdrD. These results suggested an increased electron flux in the *M. luminyensis* CZDD1 cells when cocultured with *C. malenominatum* CZB5. In addition, genes encoding proteins containing Fe-S clusters, the *brf* encoding bacterioferritin, and the cell division protein FtsZ were also markedly upregulated in coculture (Fig. 4A and Supplementary dataset S2). However, the expression of *mcrABDG* encoding the key methanogenic methyl-CoM reductase complex and the *echABCDE* encoding the membrane-bound energy-converting hydrogenase were slightly reduced in coculture. Therefore, the expression of these genes did not correlate with the higher methane production rate in the coculture. Moreover, the expressions of *echABCDE* were markedly reduced during EET methanogenesis. These suggested that during EET the energy-converting hydrogenase of *M. luminyensis* CZDD1 may not be important in energy metabolism (Fig. 4B and Supplementary dataset S2). In contrast to *M. luminyensis* B10, in which transcription of the *ech ABCDEF* was not detected(*26*), higher transcription of *echABCDE* (RS01460-RS01475, RS13925, Supplementary dataset S2) was determined in the CZDD1 H_2_-dependent culture. Consistent with the transcriptomics, quantitative RT-PCR determined a 100-fold decreased transcription of the *M. luminyensis echC* and *echF*, but about 3- and 4-fold higher transcription of *hdrD* and *fpoC* in the coculture and EET culture than in the H_2_-dependent monoculture. (Fig. 4C).

**Figure 4.**
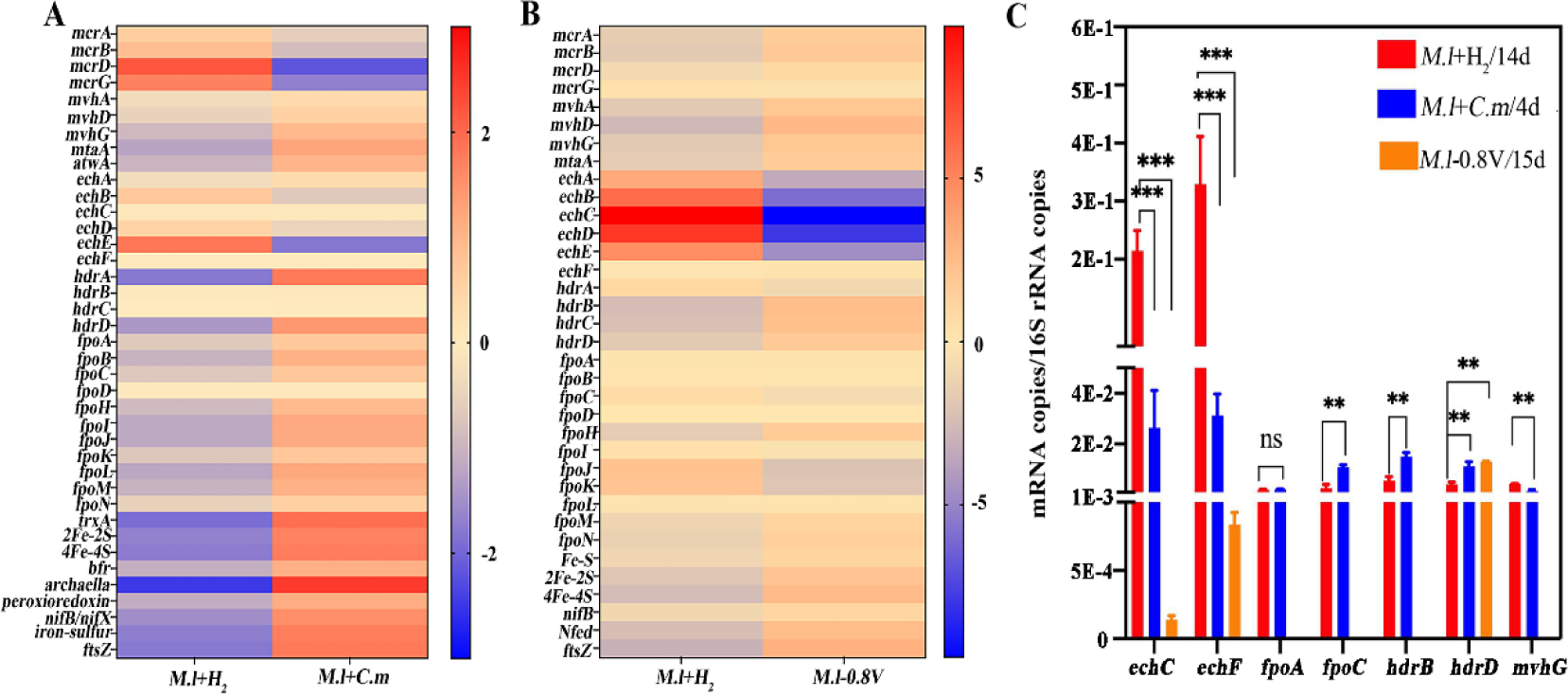
Differentially transcribed genes of *M. luminyensis* CZDD1 in H_2_-dependent versus DIET and EET growth. Heat plot representation of the differential expression (Log2) in the mid-exponential methylotrophic cultures of *M. luminyensis* grown respectively under 0.1 MPa H_2_ at day 14 (*M.l*+H_2_), and in coculture with *C. malenominatum* at day 4 (*C.m*+*M.l*) (**A**) and in a bio-electrochemical reactor at −0.8 V potential (*M.l*-0.8V) at day 4 (**B**). Blue and red represent the minima and maxima fold, respectively. (**C**) Quantitative RT-PCR of some differentially transcribed genes involved in methanogenesis in the *M. luminyensis* monoculture (red bar), coculture (blue bar) and electrochemical culture (orange bar). Three replicate experiments were performed, and the averages and standard deviations are shown in **C**. Statistics are estimated by T-test, and ***, ** and ns indicate very significant (P<0.01), significant (P<0.05) and not significant, respectively.

### The *M. luminyensis* CZDD1 pili/archaella could be involved in DIET with *C. malenominatum*

During the coculture with *C. malenominatum* CZB5, *M. luminyensis* CZDD1 had a 4-fold increase in transcription of the gene encoding the type IV pilin or archaella (Fig. 4A), and moreover this gene also ranked among the most highly expressed genes (Supplementary dataset S2). Scanning electron microscopy found pili-like appendages on the round cells of *M. luminyensis* CZDD1 and attached to the rod cells of *C. malenominatum* CZB5 (Fig. 5A). Addition of tectorigenin (TE), a chemical specifically inhibiting the expression of the type IV pilin gene (*31*), caused disappearance of the pili-like appendages of *M. luminyensis* CZDD1 (Fig. 5B). Moreover, transcription of the *M. luminyensis* CZDD1 pili gene was 2.5-fold decreased in the presence of TE (Fig. 5C insert). Importantly, methane production in the coculture was also reduced in the presence of TE (Fig. 5C). In contrast, TE did not inhibit methane production by H_2_-dependent *M. luminyensis* CZDD1 monoculture (Fig. 5D), or growth and H_2_ production of *C. malenominatum* CZB5 (Fig. 5E and 5F). Thus the *M. luminyensis* CZDD1 pili or archaella may play a role in uptake of the extracellular electrons during DIET or EET.

**Figure 5.**
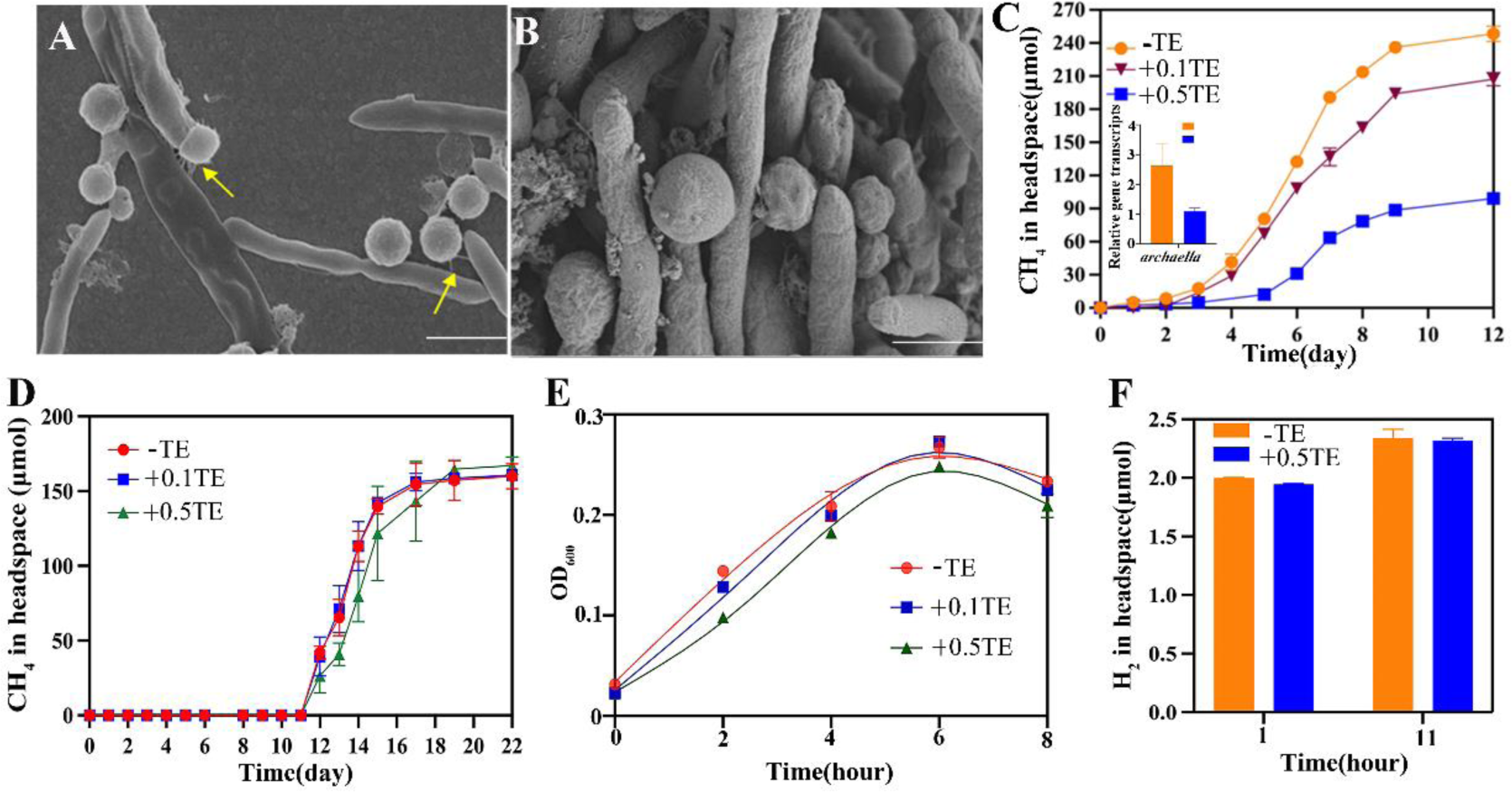
Pili/archaella-based DIET between *M. luminyensis* and *C. malenominatum*. Scanning electron microscopic images of the *M. luminyensis* and *C. malenominatum* coculture in the absence (**A**) and presence of 0.5 mM tectorigenin (**B**). Yellow arrows indicate the pili-like appendages on the spherical cells of *M. luminyensis* (**A**). Scale bars in **A** and **B** indicate 1 μm and 200 nm, respectively. Final concentrations of 0.1 mM and 0.5 mM TE were amended into the cocultures (**C**) and *M. luminyensis* monoculture (**D**), respectively. Methane productions were monitored during cultivation at 37°C. Insert in **C** shows quantitative RT-PCR determined the transcript abundance of the type Ⅳ pilin gene in a coculture amended with tectorigenin (TE) or not. Final concentrations of 0.1 mM and 0.5 mM tectorigenin were respectively amended into the *C. malenominatum* monoculture, and growth (**E**) and hydrogen production (**F**) were determined during incubation at 37°C. Triplicated experiments were performed, and the averages and standard deviations are shown.

## Discussion

Electromethanogenesis has been reported in both the aceticlastic and hydrogenotrophic methanogens. Here, we report the first DIET- and EET-driven methylotrophic methanogenesis by *M. luminyensis* CZDD1, which uses extracellular electrons from a partner bacterium or cathode to reduce methanol or the methyl group of DMAs for methane production. Archaea in Methanomassiliicoccales conserve energy exclusively from reductive methylotrophic methanogenesis, and use a different energy metabolism mode from the classical methanogens (*27*, *28*). They lack cytochromes and active energy-converting hydrogenases, but possess a membrane-bound Fpo-like complex without FpoF, an F_420_-oxidizing module. Therefore, it predicted that the Fpo-like complex of *Methanomasiliicoccus* oxidizes reduced ferredoxin and directly interacts with the membrane-associated heterodisulfide reductase subunit D (HdrD), and thus constitutes an energy-converting ferredoxin:heterodisulfide oxidoreductase complex to generate a proton motive force (PMF) (*27*). Because the methyltetrahydromethanopterin methyltransferase (Mtr) is absent, the ion gradient is believed to comprise protons and not Na^+^. Biochemical assays confirmed ferredoxin oxidation coupled heterodisulfide reduction in *M. luminyensis* B10 cell membrane preparation (*28*). Based on the physiological and transcriptomic data in this study, a putative model was proposed as shown in Figure 6, in which *M. luminyensis* might preferentially utilize extracellular electrons, possibly taken up via archaella and then transferred to the Fpo-like complex, or directly taken up via the latter. The intracellular electrons would be further transferred to membrane-associated HdrD to reduce the heterodisulfide CoM-CoB. Subsequently, the reduced CoM-SH will serve as an acceptor for methyl groups during methane production. The Fpo-like complex could also generate PMF (*28*). Therefore, *M. luminyensis* CZDD1 uses extracellular electrons to accomplish methanogenesis via an abbreviated pathway and energy conservation through a simpler electron transfer chain.

**Figure 6.**
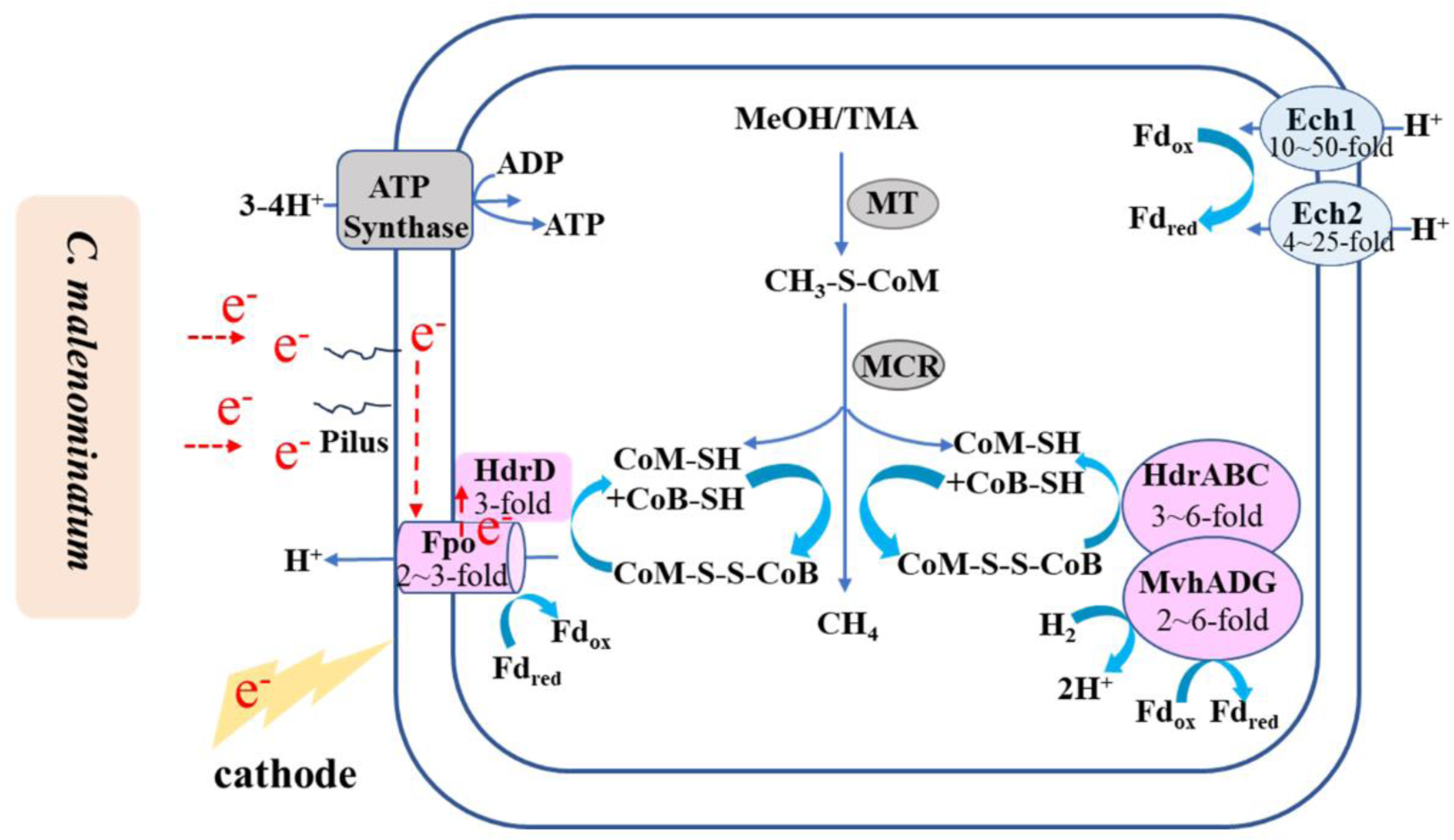
Predicted direct electromethanogenic pathway of *Methanomasilliicoccus luminyensis*. Extracellular electrons from *C. malenominatum* or cathode are accepted by the *M. luminyensis* pili and transferred to, or directed accepted by the membrane bound Fpo-like complex. Electrons are then transferred to membrane associated HdrD to reduce heterodisulfide CoM-S-S-CoB for methane production from methanol (MeOH) or trimethylamine (TMA). Protein icons shadowed in purple, blue and grey represent the encoding genes to be up- and down-regulated, and no change when *M. luminyensis* uses extracellular electron versus H_2_. MT, methylotransferase; MCR, methyl-CoM reductase.

Although the involvement of hydrogen in methane production in the coculture and the cathode reactor at −0.4 V could not be excluded due to the current inability to delete the hydrogenase genes in *M. luminyensis* CZDD1, the following experimental results verify EET but not H_2_ were used in methanogenesis, i) CO inhibition of NiFe-hydrogenases did not abolish methane production from the coculture, and ii) BES inhibition of methanogenesis reduced current consumption in the electrochemical culture. Based on the higher methane production rates and shorter lag phase observed in the co-culture and electrochemical culture compared to H_2_-culture, we propose that electron reduction of methyl compounds may represent the predominant pathway for methylotrophic methanogenesis by *M. luminyensis* CZDD1. This also conforms to the speculation of Lovley and Holmes (*6*) that free H_2_ may not be the main electron vehicle during interspecies electron transfer. Compared to other methanogens, the *Methanomassiliicoccus* strains have extremely low H_2_ threshold (*32*). They are also ubiquitous in natural environments and animal intestinal tracts (*15*), therefore could play a significant role in methane emissions via electromethanogenesis, particularly in the areas having low levels of H_2_.

DIET or EET can even activate dormant methanogenic pathways or facilitate reductive chemical reactions. The genus *Methanothrix* comprises methanogens that exclusively perform aceticlastic methanogenesis despite carrying the complete gene suite for CO_2_ reductive methanogenesis except hydrogenase. Rotaru et al. found that the electroactive bacterium *Geobacter metallireducens* enabled CO_2_ reductive methanogenesis by *Methanotrix harundinacea* (*5*). In this study, we also found that the electrochemical culture, but not the H_2_-culture, of *M. luminyensis* CZDD1 could produce methane from DMAs (Fig. 3).

In conclusion, this study reports the first EET-based methylotrophic methanogenesis by *Methanomassiliicoccus luminyensis* CZDD1, a strain isolated from DMAs contaminated paddy soil. Strain CZDD1 preferentially uses DIET or EET for methylotrophic methanogenesis, in particular from DMAs, a toxic chemical to rice. This study not only reveals a new electromethanogenic pathway, but also found that DIET and EET can activate the inactive methanogenic pathways harbored in methanogens. Based on the ubiquitous distribution of *Methanomassiliicoccus* and methylotrophic methanogenic pathways in natural environments, electromethanogenesis including electron-driven methylotrophic methanogenesis could contribute significantly to global methane emission.

## Materials and Methods

### Strains and cultivations

Pre-reduced and modified MM medium (*25*) was used as the basic medium, which includes 0.5g KH_2_PO_4_, 0.4g MgSO_4_·7H_2_O, 5g NaCl, 1 g NH_4_Cl, 0.05g CaCl_2_ 2H_2_O, 1.6g sodium acetate, 1g yeast extract, 1g tryptone and 4g NaHCO_3_. To culture *C. malenominatum* CZB5, 0.1 MPa N_2_ was added in the headspace, while culturing *M. luminyensis* CZDD1, 15 mM methanol and 0.1 MPa H_2_ in the headspace were used. The coculture of strain CZDD1 and CZB5 were grown in modified MM medium amended with 15 mM methanol and 0.1 MPa N_2_. *Geobacter metallireducens* ATCC53774 and its pili defect mutant *ΔpilA* were routinely grown in the ferric citrate medium amended with 20 mM ethanol (*2*). Cocultures of *M. luminyensis* with *G. metallireducens* wild type or *Δ pilA* strain were initiated with by 10% inoculation (0.1 mg total cell protein) of each strain into the same modified MM medium containing 20 mM ethanol and 40 mM methanol and under 0.1MPa N_2_ phase. To assay carbon monoxide inhibition on methane production, 0.2 MPa mixture of 10% CO and 90% N_2_ was pulsed into the co-culture and monoculture. All the cultures were included at 37`°`C.

### Characterization of the bio-electrochemical cultures

Chronoamperometry was carried out using a Bio-logic science instrument potentiostat CHI 1000C (CH Instrument, TX). Electrochemical experiments were set up in the H cell reactor, in which 150 mL-anode and cathode chambers were separated by a Nafion 117 proton-exchange membrane (Dupond, USA), and equipped with an Ag/AgCl reference electrode (CH Instruments), a Pt wire counter electrode (Alfa Aesar), and a 6.35-mm-thick graphite felt working electrode with a 16-mm radius (Alfa Aesar), in addition of two sampling ports. For onset of the electrochemical experiment, H cell reactor was first autoclaved and then flushed with 100% N_2_ gas for no less than 15 min to achieve an oxygen-free headspace. Next, the anode and cathode compartments were filled with 80 mL sterilized and pre-reduced modified MM medium omitting sulfide, and resazurin inside an anaerobic box (Thermofisher). The H-cell reactors were continuously purged with N_2_ gas to maintain anaerobic condition. The mid-exponential culture of *C. malenominatum* CZB5 was 10% inoculated into the MM medium within the cathode chamber with an applied cathode potential set at +0.4 V versus Ag/AgCl. The mid-exponential culture of *M. luminyensis* CZDD1 was 10% inoculated in MM medium containing 15 mM methanol in cathode chamber under the cathode potential sequentially set at −0.4 V, −0.6 V and −0.8 V. The electrochemical culture was mixed with a magnetic bar during incubation at 37`°`C. H cell reactors were connected to potential station, and currents are reported as a function of the geometric surface area of the electrode.

### Total cell protein determination

The *M. luminyensis* CZDD1 cells for inoculation were first quantified as the total cell protein. Cell pellets were resuspended with 1 mL PBS buffer and grinded in liquid nitrogen. After centrifugation at 5000 g at 4 °C for 15 min. the supernatant was used to determine total cell protein using the BCA Protein Assay Kit (Thermo Fisher Scientific, Germany).

### Ferrozine assay of ferric iron reductive activity of *C. malenominatum*

Ferric citrate at the final concentrations of 2.5 mM, 5 mM, and 7.5 mM was supplemented into the *C. malenominatum* CZB5 culture at the late-log phase, and incubated at 37 °C. 0.1% phenanthroline was added to samples of the culture and the absorbance was measured spectrophotometrically at 562 nm according to Light, et al. (2018)(*8*). Ferrous iron concentration standard curve was generated using FeCl_2_. Triplicate experiments were performed.

### Measurements of methane, hydrogen, and ^13^C/^12^C ratios of methane and carbon dioxide

Methane was measured using GC-14B gas chromatograph (Shimadzu, Japan) equipped with flame ionization detector and C18 column as described previously (*33*). The temperature parameters were used as follows: column at 50 °C, injector at 80 °C, and detector at 130 °C. Hydrogen was measured using GC-4100 (GWAI) equipped with a TCD detector and column temperature parameters were used as follows: column at 80 °C, injector at 150 °C and detector at 90 °C. In addition, ^13^C-labeled DMAs(V) was synthesized according to Chen et al. (*34*). The ^13^C/^12^C isotope ratios of methane and carbon dioxide in headspace were determined using GC - IRMS (isotope ratio mass spectroscopy, Thermo Fisher Scientific, Germany).

### Fluorescent in situ hybridization

The mid-exponential phase of the co-culture of *M. luminyensis* CZDD1 and *C. malenominatum* CZB5 was collected by centrifugation at 3500 g for 10 min. Cell pellets were rinsed with PBS, fixed in 4% paraformaldehyde for 12 h at 4℃, and then dehydrated with gradient concentrations of ethanol from 50 to 100%. Fluorescent in situ hybridization (FISH) was carried out in a 20 mM Tris buffer (pH 7.4) containing 900 mM NaCl, 0.01% SDS (w/v), and 40% formamide (v/v) at 46℃ for 2 h. Probes used were 5’-[CY3] GCTGCCTCCCGTAGGAGT-3’ targeting the *C. malenominatum* 16S rRNA and 5’-[CY5] GTGCTCCCCCGCCAATTCCT-3’ targeting the *M. luminyensis* 16S rRNA. Fluorescent cells were observed under LSM 800 confocal microscope (Leica Microsystems, Buffalo Grove, IL, USA).

### Scanning electron microscopy

The mid-exponential cells from the co-culture of *M. luminyensis* CZDD1 and *C. malenominatum* CZB5 without or with tectorigenin (TE) were collected by centrifugation at 3500 g for 10 min. Cell pellets were rinsed with sterile phosphate-buffered saline (PBS), and fixed overnight at 4℃ in 0.1 M phosphate buffer (pH 7.0) containing 2.5% glutaraldehyde. Fixed cells were then dehydrated by gentle agitation in gradient concentrations of ethanol from 30 to 100%. After dried at 55℃, cells were observed under a Quanta 200 scanning electron microscope (FEI, Hillsboro, OR, USA).

### RNA library preparation and Illumina Hiseq sequencing

The mid-exponential cells were collected from *M. luminyensis* CZDD1 H_2_-dependent culture, its co-culture with *C. malenominatum* CZB5 and the electrochemical culture with cathode potential set at −0.8 V. Total RNA was extracted using the RNAiso Plus kit (Takara, Dalian, China) according to the manufacturer’s instructions. RNA quality was determined using 2100 Bioanalyser (Agilent) and quantified using the ND-2000 (NanoDrop Technologies).

High-quality RNA sample (OD260/280=1.8∼2.2, OD260/230≥2.0, RIN≥6.5, 23S:16S≥1.0,>10μg) is used to construct sequencing library.

Total RNA of 5 μg was used to construct strand-specific RNA-sequencing libraries using the TruSeq RNA sample preparation Kit from Illumina (San Diego, CA). Briefly, rRNA was removed by the RiboZero rRNA removal kit (Epicenter), and then RNA was fragmented using fragmentation buffer. cDNA synthesis, end repair, A-base addition and ligation of the Illumina-indexed adaptors were performed according to Illumina’s protocol. Libraries were then size selected for cDNA target fragments of 200–300 bp on 2% Low Range Ultra Agarose followed by PCR amplified using Phusion DNA polymerase (NEB) for 15 PCR cycles. After quantified by TBS380, paired-end libraries were sequenced on a Illumina NovaSeq 6000 (150bp*2, Shanghai BIOZERON Co., Ltd).

### Reads quality control and mapping

The raw paired end reads were trimmed and quality controlled by Trimmomatic with parameters (SLIDINGWINDOW:4:15MINLEN:75) (version0.36 http://www.Usadellab.org/cms/uploads/supplementary/Trimmomatic). The clean reads were aligned to the reference genome with orientation mode using Rockhopper (http://cs.wellesley.edu/~btjaden/Rockhopper/) software. Rockhopper was a comprehensive and user-friendly system for computational analysis of bacterial RNA-seq data. As input, Rockhopper takes RNA sequencing reads generated by high-throughput sequencing technology. This software was used to calculate gene expression levels with default parameters.

### Differential expression analysis and function enrichment

To identify DEGs (differential expression genes) between two samples, transcript abundances of genes were calculated based on Fragments Per Kilobase of read per Million mapped reads (RPKM). EdgeR (https://bioconductor.org/packages/release/bioc/html/ edgeR. html) was used for differential expression analysis. DEGs between two samples were selected based on the following criteria: the logarithmic fold change was greater than 2, and the false discovery rate (FDR) was less than 0.05. To understand the functions of the differential expressed genes, GO functional enrichment and KEGG pathway analysis were carried out by Goatools (https://github.com/tanghaibao/Goatools) and KOBAS (http://kobas.cbi.pku.edu.cn/kobas3) respectively. DEGs were significantly enriched in GO terms and metabolic pathways when their Bonferroni-corrected P-value was less than 0.05.

### Quantitative reverse transcription PCR (RT-qPCR)

RNA extract used for sequencing above was treated with RNase-Free DNase (Takara) to remove contaminated DNA, and cDNA was synthesized using the M-MLV Reverse Transcriptase cDNA Synthesis Kit (Takara) following the manufacturer’s instructions. Using the primers listed in Supplementary Table S1. Quantified PCR was performed using SYBR Premix Ex Taq Kit (Takara Bio Inc., Japan), and carried out on ABI Prism 7000 sequence detection system (Applied Biosystems USA). Each qPCR mixture contained 12.5 μL SYBR qPCR mix (TOYOBO), 5 μL cDNA, 100 nM of each primer, and double-distilled H_2_O to a final volume of 25 μL. PCR was initiated at 95 °C for denaturation for 30 s and 35 cycles as follows: denature at 95 °C for 10 s, annealing at the Tm listed in Supplementary Table S1 for 30 s, and elongation at 72 °C for 30 s. Fluorescence data were collected during elongation. To estimate mRNA copies of the tested genes, a standard curve of each gene was generated by quantitative PCR on 10-fold serially diluted plasmid DNA cloned with the gene. 16S rRNA gene was used as the biomass reference. Triplicate samples were determined and each measurement was repeated at least three times.

## Data availability

The genomic sequences of and transcriptomic data of the *C. malenominatum* CZB5 and *M. luminyensis* CZDD1 have been deposited in NCBI SRA database under the accession numbers of PRJNA1032237 and PRJNA1028373, respectively.

## Acknowledgements

We thank Professor Yahai Lu for talking about electrochemical experiment and CO inhibition, and Mr Yuhong Zhong for providing strain *G. metallireducens.* This work was supported by National Natural Science Foundation of China (92251302, 91851211 and 32070061), and the Second Tibetan Plateau Scientific Expedition and Research (STEP) Program (2019QZKK0304).

