## supplemental materials for "Extracellular electron transfer drives efficient H_2_-independent methylotrophic methanogenesis by *Methanomassiliicoccus,* a seventh order methanogen"

**This file contains:**

Supplementary Tables S1.

Supplementary Figure S1-S3.

Caption for supplementary dataset S1: Differential transcriptome of *C. malenominatum* CZB5 in its monoculture vs in the coculture with *M. luminyensis* CZDD1

Caption for supplementary dataset S2: Differential transcriptomes of *M. luminyensis* CZDD1 in coculture with *C. malenominatum* CZB5, and in electrochemical culture vs. its H_2_-dependent methylotrophic culture, respectively

**Table S1.** Primers used in this study.

| Primers | Sequences (5’-3’) | Tm (℃) | Purpose |
| --- | --- | --- | --- |
| *mvhG*-F | AACCGACCACCAAGTCCG | 57 | PCR |
| *mvhG*-R | CTGCCTTGAGGGGCATCT |  |  |
| *mvhG*-qF | GCACCGTTCCTATGCTCAGA | 57 | qPCR |
| *mvhG*-qR | GTCGCCGTCGTAGAAGTTGA |  |  |
| *hdrB*-F | AAGTACGCGCTGTTCCTG | 55 | PCR |
| *hdrB*-R | CAGCTGGGACAGGTGGAT |  |  |
| *hdrB*-qF | GGCATTGAGTCCTCTACCCG | 63 | qPCR |
| *hdrB*-qR | CGTCGAACAGAGAGCCGTAG |  |  |
| *fpoA*-F | AGCATGATGTCGATCGTG | 58 | PCR |
| *fpoA*-R | GCCTATCGGCTGAGGT |  |  |
| *fpoA*-qF | GACACGGTCCCTTACCCTG | 58 | qPCR |
| *fpoA*-qR | CTCTATGGTGGCGGCCTTC |  |  |
| *fpoC-*F | AGCGTACAACTCACCGTGG | 55 | PCR |
| *fpoC-*R | CCCTACGGTGTAGTCCTTCC |  |  |
| *fpoC*-qF | AAGCTGTGCACCTTCCTC | 63  59 | qPCR |
| *fpoC-*qR | TGTCGTAGGTCTCCCTCT |  |  |
| *echC*-F | ATCACAAGCAGAGAAGCC |  | PCR |
| *echC*-R | GTCTCGTTACGTTCCCT |  |  |
| *echC*-qF | GCATGCTCAACAAGGGCAAC | 57 | qPCR |
| *echC*-qR | CCGTCGATCAATGCCTCTGG |  |  |
| *echF*-F | ATGAGCGAACTGCTAACG | 59 | PCR |
| *echF*-R | CGTCATGGCCAGGAGCAT |  |  |
| *echF*-qF | CCTCGCCTATTCTACCGTCG | 59 | qPCR |
| *echF-*qR | AACACTCCCACGAAGATGCC |  |  |
| *hdrD*-F | CCGGAAATCAACAGAGAG | 52 | PCR |
| *hdrD*-R | GAAGCGGAGGCCTTCTCT |  |  |
| *hdrD*-qF | CAAGTACTTCGCCAACCCCT | 61 | qPCR |
| *hdrD*-qR | CCCCGGGGATCTTCTTGATG |  |  |
| *Archaella*-F | CAGTGTCCCCTGTGATTGCC | 55 | PCR |
| *Archaella*-R | GAACGTGCAAGCCGCTA |  |  |
| *Archaella*-qF | TTACAAGGTCGGGGTAGGCT | 60 | qPCR |
| *Archaella*-qR | GTCGTTTCCGTCAACGTCCT |  |  |

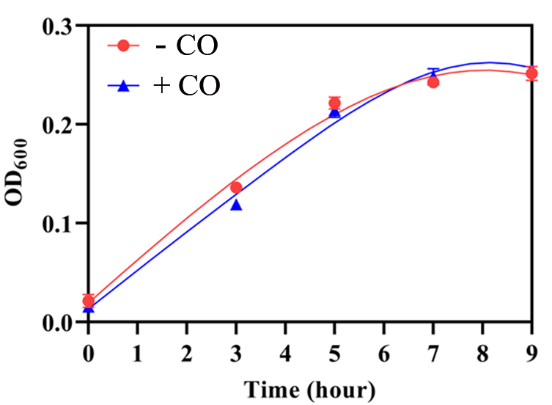

**Figure S1. Effect of carbon monooxide on** ***C. malenominatum* growth.** A gas mixture 10% CO (+) and 90% N_2_ at 0.2 MPa was pulsed to the modified MM culture of *C. malenominatum* CZB5, and culture without pulsed CO (-) was included as a control. Optical density at 600 nm (OD_600_) was monitored during incubation at 37ºC. Three independent experiments were performed, and the averages and standard deviations are shown.

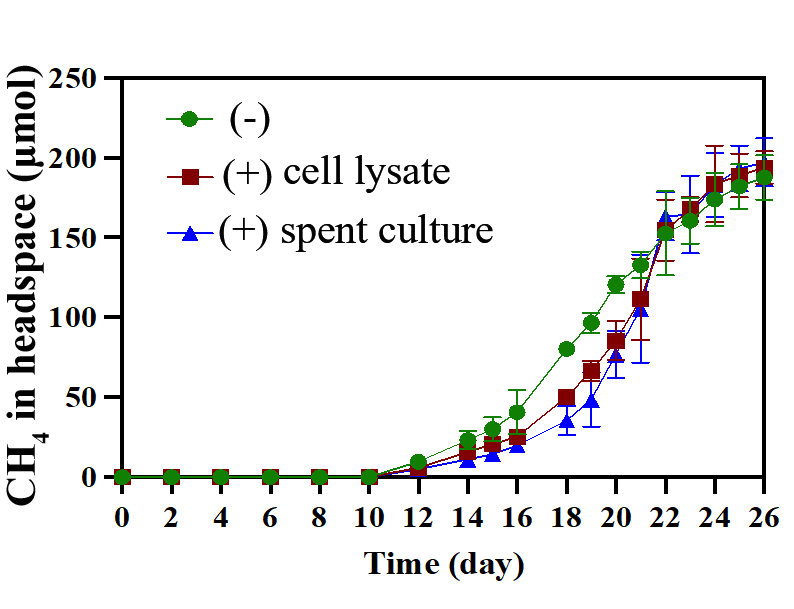

**Figure S2. Effects of** **the *C. malenominatum* CZB5 cell components or products on enhancing methylotrophic methanogenesis by *M. luminyensis*.** The exponential culture of strain CZB5 grown in modified MM medium was centrifuged, and the supernatant was used as the spent culture and the cell pellet was sonicated to prepare cell lysate, respectively. Spent culture of 25 ml and cell lysate of 10 mg total protein were respectively added to the H_2_ + methanol culture of *M. luminyensis*, and no amendment was used as control (-). Methane in headspace was monitored during incubation at 37 ºC. Three independent experiments were performed, and the averages and standard deviations are shown.

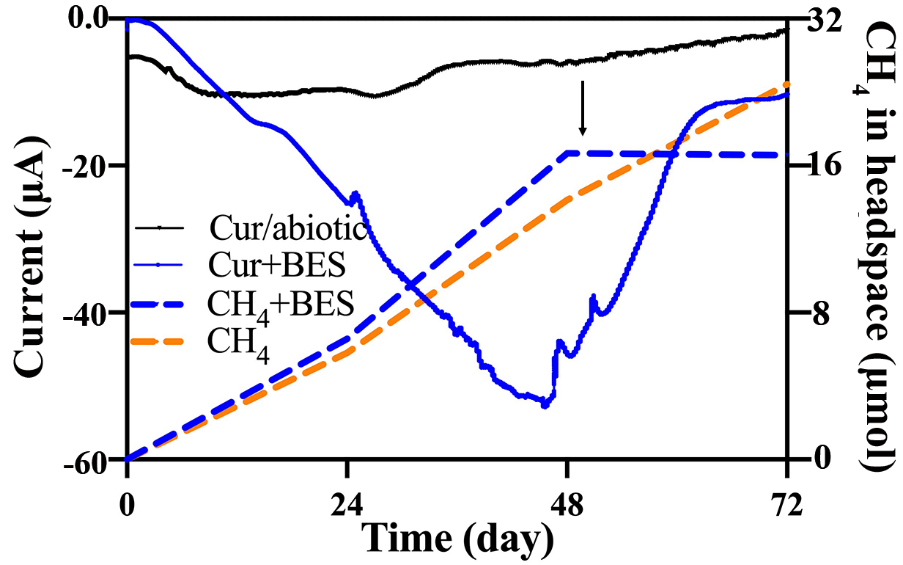

**Figure S3.** **The activity of *M. luminyensis* B10 utilizing cathodic electrons for methanogenesis.** *M. luminyensis* B10 was 10% inoculated into the modified MM medium containing 400 μmol methanol in a H-cell electrochemical reactor with cathode potential at -0.4 V, and continuously purged with 0.1 MPa N_2_. A H-cell reactor without inoculation was included as the abiotic control. A final concentration of 10 mM 2-Bromoethanesulfonate (BES) was added 48 h post inoculation, and methane production coupled current consumption was monitored. Two independent experiments were performed and one representative was shown.
